# Dynamic Hippo pathway activity underlies mesenchymal differentiation during lung alveolar morphogenesis

**DOI:** 10.1101/2023.10.17.561252

**Authors:** Fatima N. Chaudhry, Nigel S. Michki, Dain L. Shirmer, Sharon Mcgrath-Morrow, Lisa R. Young, David B. Frank, Jarod A. Zepp

**Affiliations:** Division of Pulmonary and Sleep Medicine, Department of Pediatrics, Children’s Hospital of Philadelphia. Philadelphia, Pennsylvania, USA, 19104; Division of Cardiology, Department of Pediatrics, Children’s Hospital of Philadelphia. Philadelphia, Pennsylvania, USA, 19104

## Abstract

Alveologenesis, the final stage in lung development, substantially remodels the distal lung, expanding the alveolar surface area for efficient gas exchange. Secondary crest myofibroblasts (SCMF) exist transiently in the neonatal distal lung and are critical for alveologenesis. However, the pathways that regulate SCMF function, proliferation, and temporal identity remain poorly understood. To address this, we purified SCMFs from reporter mice, performed bulk RNA-sequencing, and found dynamic changes in Hippo-signaling components during alveologenesis. We deleted Hippo effectors, Yap/Taz, from Acta2-expressing SCMFs at the onset of alveologenesis, causing a significant arrest in alveolar development. Using scRNA-seq, we identified a distinct cluster of cells in mutant lungs with altered expression of marker genes associated with proximal mesenchymal cell types, airway smooth muscle (ASM), and alveolar duct myofibroblasts (DMF). Using lineage tracing, we show that neonatal Acta2-expressing SCMFs give rise to adult DMFs and that Yap/Taz mutants have an increase of persisting DMF-like cells in the alveolar ducts. Our findings identify plasticity in neonatal lung myofibroblasts and demonstrate that Yap/Taz are critical for maintaining lineage commitment along the proximal-distal axis.

## Introduction

The human lung is estimated to have more than 300 million alveoli that cover approximately 75 m^2^ of surface area, facilitating efficient gas exchange^1^. Alveolar morphogenesis and maturation occur immediately before and after birth through a developmental process called alveologenesis. During this phase, coordinated activities of many cell types sculpt and multiply the alveoli. Infants born prematurely have disrupted alveologenesis and are at risk for developing neonatal lung diseases, often leading to lifelong complications^1–3^.

Epithelial and mesenchymal cell types are critical for the expansion and maturation of the alveoli. Alveolar type 2 (AT2) epithelial cells proliferate and increase surfactant production, while AT1 cells flatten and spread out to accommodate the demands of a growing surface area^4, 5^. The mesenchyme, including fibroblasts and smooth muscle, is highly heterogeneous and is critical for alveologenesis. Pdgfra-expressing mesenchymal cells proliferate, produce extracellular matrix (ECM), and support the expansion of AT2 cells by secreting growth factors^6–9^. A unique neonatal Pdgfra-expressing myofibroblast called the secondary crest myofibroblast (SCMF) exists transiently during lung alveologenesis and exerts physical force to help shape new alveoli^10–13^.

Previously, we showed that the SCMF is a transcriptionally and functionally distinct mesenchymal cell lineage^13^. After birth, SCMFs expand in number and exert contractile forces in the alveolar interstitium. SCMFs, but not other Pdgfra-expressing fibroblasts, exert more tractional forces to generate an expansive network of alveolar septa^13^. Because of its contractile properties and unique transient existence, uncovering signaling pathways regulating this cell type could inform strategies for treating neonatal lung diseases through enhanced alveolar maturation.

Here, we studied how the Hippo signaling pathway impacts the function and differentiation of SCMFs. Our data demonstrate the dynamism of YAP in SCMFs throughout alveologenesis. *In vivo*, conditional knockout of Yap/Taz from Acta2-expressing SCMFs during alveologenesis is detrimental to alveolar maturation. We identify persistent cells in the mutant lungs residing in the alveolar ducts, a transitional zone between the airway and alveoli. These observations highlight the cell-type-specific role of Hippo pathway signaling in controlling neonatal myofibroblast development and suggest that biomechanical forces influence the allocation of distinct mesenchymal lineages.

## Results

### RNA-seq analysis of SCMFs identifies Hippo signaling dynamics during alveologenesis

Transient neonatal myofibroblasts, or SCMFs, are critical for proper alveolar maturation. Our previous work demonstrated that we can purify SCMFs for functional assays, using murine reporter lines^13^. To identify novel signaling pathways that may contribute to SCMF function or differentiation, we performed bulk RNA sequencing from purified SCMFs during early alveologenesis. Using *Acta^DsRed^:Pdgfra^GFP^* dual reporter mice, we profiled the RNA from DsRed/GFP double-positive cells spanning the perinatal period from E17.5 to P10 when SCMFs are abundant (**Fig. 1A**). Principle component analysis indicated that changes in gene expression were primarily attributed to the transition from embryonic to postnatal timepoints (PC1, 21.31% variance) and secondarily to transitions during the period of postnatal lung development (PC2, 16.51% variance) (**Fig. 1B**). Analyzing these data for hallmark signaling pathways that have been implicated in lung mesoderm development, we found that SHH, PDGF, and WNT signaling pathways are highest at the embryonic time point, E17.5, and then decline postnatally (**Fig. 1C**). The Hippo signaling pathway, critical for determining cellular growth in the context of mechanical cues, showed increasing activity in the SCMF (**Fig. 1C**). The genes that were enriched in the Reactome database showed dynamic expression between embryonic and postnatal SCMFs. Many were known downstream target genes of TEA domain (TEAD) transcription factors. Notably, genes encoding Hippo pathway effectors, *Lats1/2*, *Mob1a/b*, *Sav1*, *Yap1*, and *Wwtr1* (Taz), consistently had higher expression at the P10 timepoint (**Fig. 1D**) ^14^. After the SCMF cell numbers peak at P7, a rapid decline through apoptosis follows^13, 15^. In line with this was increased *Casp3* expression in the P3 and P7 samples (**Fig. 1D**). Together, these data implicate active and dynamic Hippo signaling pathway activity in the SCMF that may mediate SCMF function and apoptosis.

**Figure 1.**
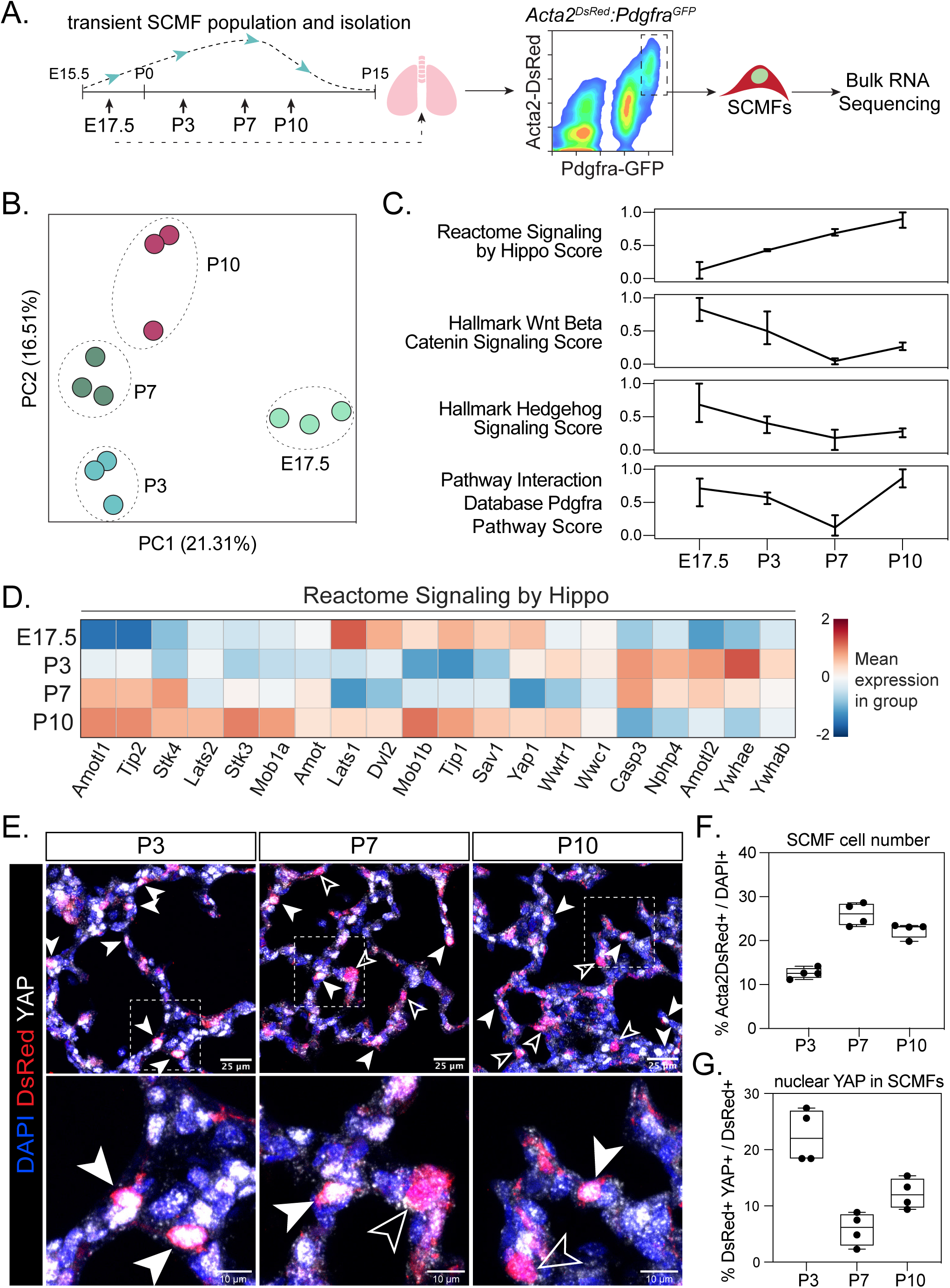
Secondary crest myofibroblasts (SCMF) have enriched Hippo signaling activity. (A) Experimental schematic for isolation of SCMF from *Acta2^DsRed^:Pdgfra^H2B:EGFP^* mice at the indicated time points followed by bulk RNA-seq. (B) Principal component analysis (PCA) plots of the bulk RNA-seq. Dots represent biological replicates grouped by the indicated time point. (C) Signaling scores of pathways with known relevance to early lung development. (D) Heatmap showing expression of genes associated with Reactome Signaling by Hippo. (E) Representative imaging results of immunostaining for YAP at the indicated time points on tissue from *Acta2^DsRed^* mice. Scale bars, 25 µm. Lower, high magnification of alveolar regions showing DsRed+ fibroblasts with (solid arrows) or without (empty arrows) nuclear YAP. Scale bars, 10 µm. (F) Quantification of Acta2DsRed+ SCMFs. (G) Quantification of nuclear YAP+ DsRed+ SCMFs. Bulk RNA-seq from n=3 mice per time point, YAP imaging and quantification are from n=4 mice, presented as mean ± SD.

To assess Hippo pathway activity in SCMFs, we examined lung tissue from *Acta2^DsRed^* reporter mice for subcellular localization of YAP protein (**Fig. 1E**). When the Hippo pathway is “on”, YAP/TAZ is actively phosphorylated and resides in the cytoplasm, marked for proteasomal degradation^14^. When the Hippo pathway is “off,” the upstream kinases are inhibited and YAP/TAZ can then translocate to the nucleus, acting as cofactors for the TEAD family of transcription factors to regulate target gene expression^14^. While YAP nuclear-cytoplasmic shuttling is dynamic, at P3, we found that 20-30% of SCMFs (alveolar DsRed+ cells) have nuclear localized YAP protein (**Fig. 1F, G**). Notably, the percentage of SCMFs with nuclear YAP declines at P7 and P10 when SCMF numbers peak (**Fig. 1F, G**). Together, these data indicate that Hippo pathway has dynamic activity in the SCMF. Further, nuclear translocation of YAP inversely correlates with SCMF numbers.

### Deletion of Yap/Taz in myofibroblasts results in alveolar simplification

To determine the role of Yap/Taz in SCMFs, we crossed a tamoxifen-inducible Cre-recombinase, *Acta2^CreERT2^*, with *Yap^FF^/Taz^FF^* alleles to generate a conditional knockout model. We’ve shown that *Acta2^CreERT2^*(αSMA) labels airway and vascular smooth muscle and is highly specific for SCMFs during alveologenesis, mostly sparing other fibroblasts^13^. We induced Cre activity with tamoxifen at postnatal day 1 (P1) in *Acta2^CreERT2^:Yap^FF^:Taz^FF^:R26R^eYFP^*knockout (KO) and heterozygous littermate control mice (Ctrl), and harvested lungs at P7 to assess alveolar maturation and SCMF abundance (**Fig. 2A**). The lung tissue from KO mice showed reduced numbers of septa and alveolar simplification at P7 (**Fig. 2B**). Elastin increases during alveologenesis and is thought to contribute to septation^1, 3, 16^. Therefore, we assessed elastin fiber organization as indicated by hydrazide staining between Ctrl and KO and observed no differences in hydrazide intensity (**Fig. 2C**) ^17^. We next used a three-dimensional approach to examine alveolar morphometry in precision-cut lung slices (PCLS). The alveolar diameters were significantly higher in the KO (**Fig. 2D**). To determine if Yap/Taz deletion results in a loss of SCMF differentiation, we stained Ctrl and KO tissue with SM22α (TAGLN), a marker for SCMFs. We found no difference in the percentage of SM22+ cells in the alveoli of KO tissue (**Fig. 2E, F**). Heterozygous mice and single gene knockout animals of Yap or Taz did not have obvious alveolar simplification at P7 (**Fig. S1A**). These data demonstrate that loss of the downstream effectors of the Hippo pathway, Yap/Taz, in *Acta2+* cells leads to functional impairment and alveolar simplification.

**Figure 2.**
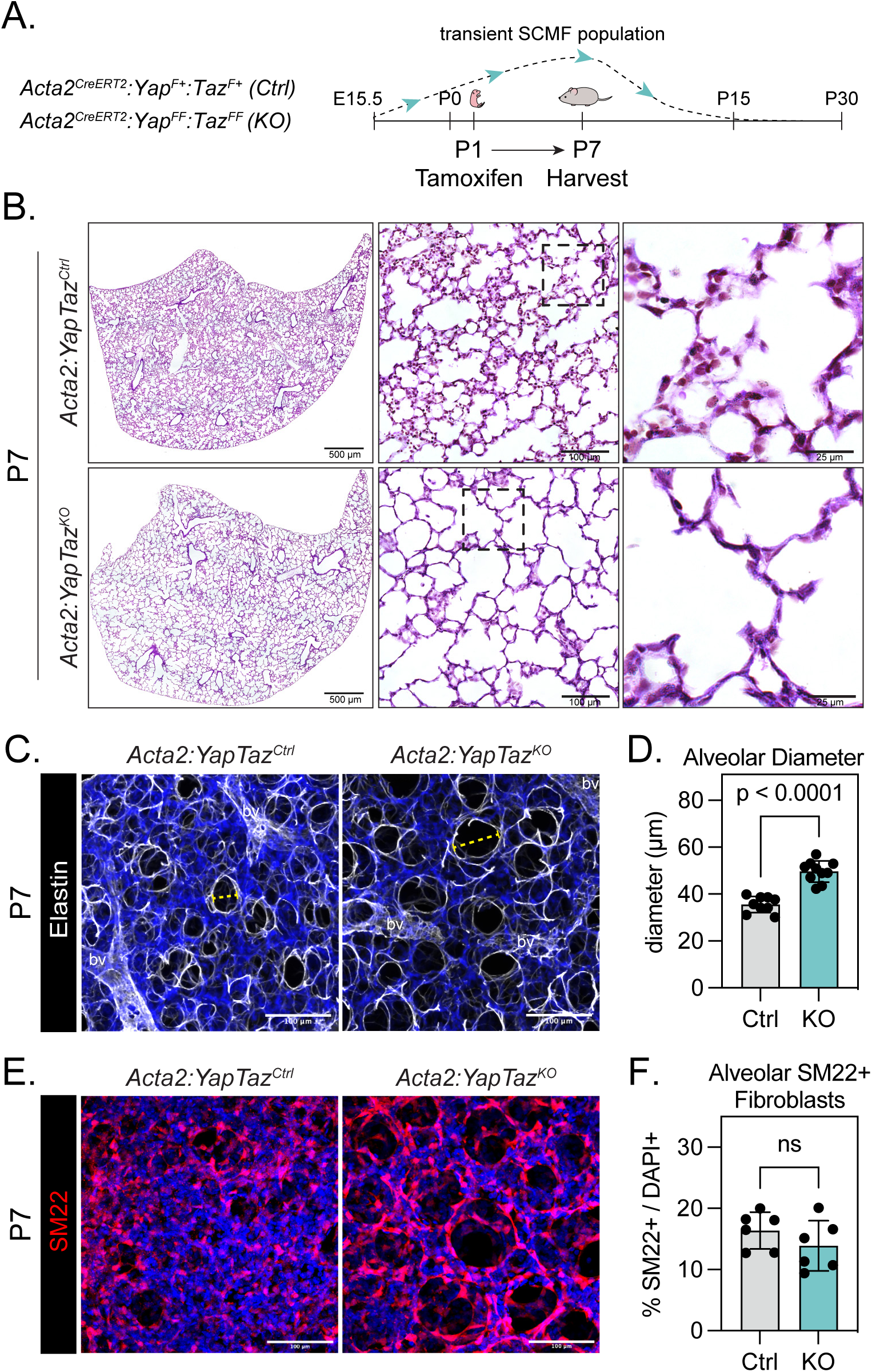
Yap/Taz deficient SCMFs attenuate alveolar development during early alveologenesis. (A) Experimental design: Recombination was induced in pups at P1, and lungs were analyzed at P7 during peak SCMF abundance. (B) Representative H&E images of Ctrl and KO lungs. Scale bars (left to right), 500 µm, 100 µm, 25 µm. (D) PCLS images of Ctrl and KO lung sections highlighting elastin fibers around alveoli using fluorescent hydrazide. The yellow dashed line denotes an alveolar diameter measurement quantified in (D), aw, airway; bv, blood vessel. Scale bar, 100 µm. (D) Quantification of alveolar diameters shows a marked increase in KO mice. Ctrl n=9, KO n=10. (E) PCLS immunostaining for SM22 in Ctrl and KO lung sections. Scale bar, 100 µm. (F) No significant change in the percentage of SM22-positive SCMFs. Ctrl n=6, KO n=6. Data presented as mean ± SD.

We aged Ctrl and KO mice to three weeks (P23) when SCMFs are normally no longer present and alveologenesis is nearly complete (**Fig. 3A**). The effects of Yap/Taz deletion in the *Acta2*-expressing cells resulted in simplification of alveoli as shown by H&E sections, elastin staining, and alveolar diameter measurements (**Fig. 3B-D**). At P23, the SM22 staining in the alveolar region had resolved entirely in Ctrl and KO tissue (**Fig. 3E, F**). Single gene knockout animals of Yap or Taz exhibited normal alveolar development at P23 (**Fig. S1B**). Due to the alveolar simplification defect observed in the mutant lungs, we next tested whether epithelial and endothelial numbers were normal. We did not observe significant changes in AT1, AT2, or endothelial cell numbers (**Fig. S2**).

**Figure 3.**
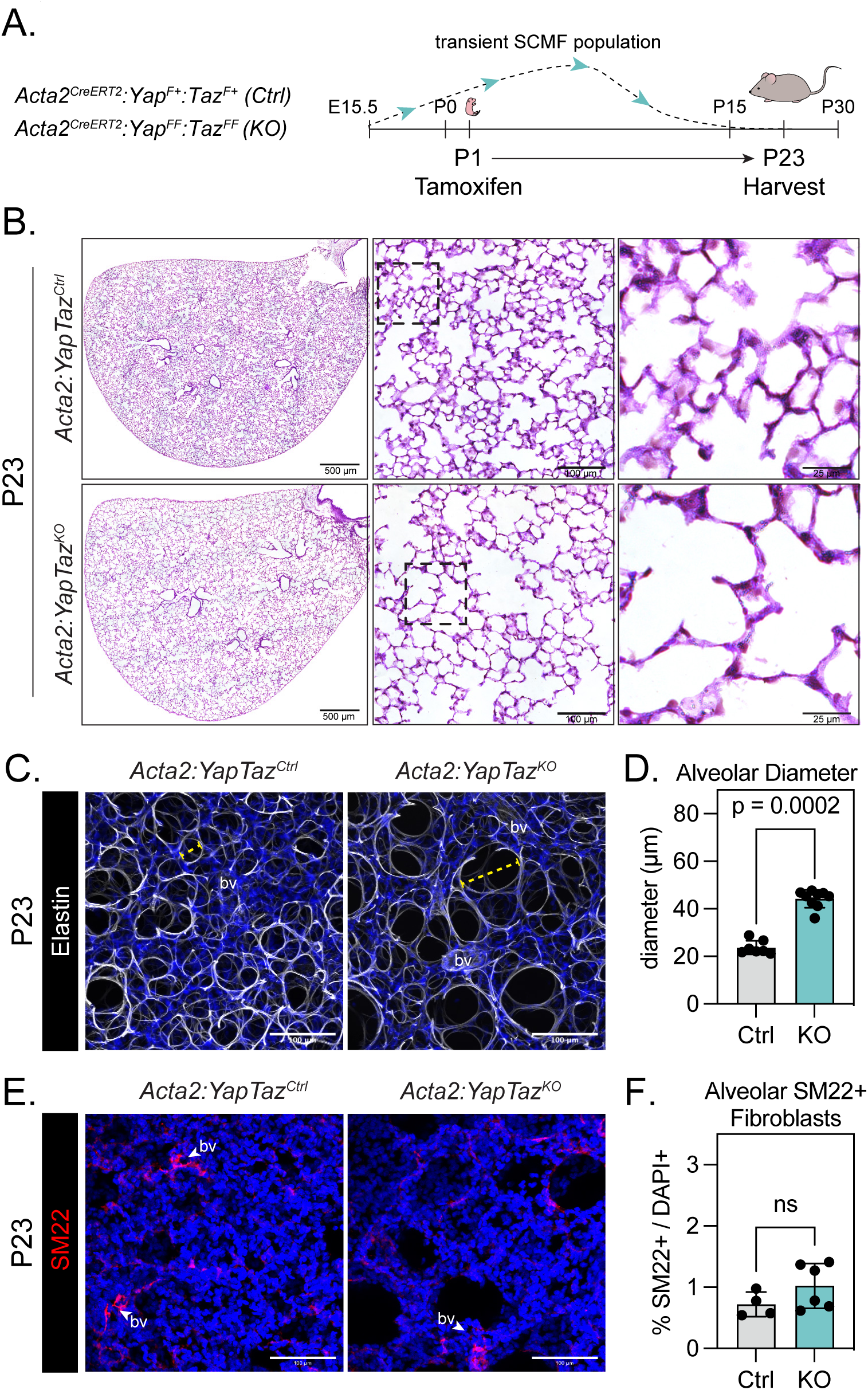
Impaired alveolarization in P23 KO lungs. (A) Experimental design: Ctrl and KO lungs were analyzed at P23 after SCMFs undergo developmental apoptosis. (B) Representative H&E images of Ctrl and KO lungs. Scale bars (left to right), 500 µm, 100 µm, 25 µm. (C) Elastin staining of Ctrl and KO PCLS. aw, airway; bv, blood vessel. Scale bar, 100 µm. (D) Quantification of alveolar diameters shows a significant increase in KO lungs, as exemplified by the dotted line in (C). Ctrl n=7, KO n=9. (E) PCLS immunostaining of Ctrl and KO lung sections stained for SM22. Scale bar, 100 µm. (F) SM22+ fibroblast population resolves in both Ctrl and KO lungs at P23. Ctrl n=4, KO n=6. Data presented as mean ± SD.

### Single-cell RNA sequencing uncovers Hippo-pathway-mediated gene expression changes

Because of the pronounced alveolar simplification in our conditional knockout model and to better understand how Yap and Taz control SCMF function, we performed single-cell RNA sequencing (scRNA-seq) of lungs from Ctrl and KO mice at P7 and P23 (**Fig. 4A**). We captured transcriptional profiles from the expected cell types in epithelium, endothelium, immune, and mesenchyme (**Fig. 4B**). Next, we sub-sampled clusters enriched in mesenchymal marker genes. The UMAP projections and re-clustering showed that we captured all previously described subsets of mesenchymal cells (**Fig. 4C, D**). While naming conventions vary, recent mesenchymal scRNA-seq data sets consistently capture two types of *Pdgfra*-expressing fibroblasts. These cells are also called *Col14a1^+^*/mesenchymal alveolar niche cells (MANC)/alveolar fibroblast 2 (AF2) or *Col13a1^+^*/lipofibroblasts/AF1^8, 18–20^. For simplicity, we refer to these *Pdgfra*-expressing fibroblasts as either adventitial (adv. fib.) or alveolar (alv. fib.), respectively. The *Pdgfrb*-expressing cell types included pericytes (*Pdgfrb^+^Acta2^low^*) and vascular smooth muscle (VSM). In addition to the VSM, we also captured several *Acta2* expressing cell-types, including the airway smooth muscle (ASM), alveolar ductal myofibroblasts (DMF), and SCMF (**Fig. 4C**). Marker gene analysis showed distinct and high expression in the given mesenchymal subtype (**Fig. 4D**).

**Figure 4.**
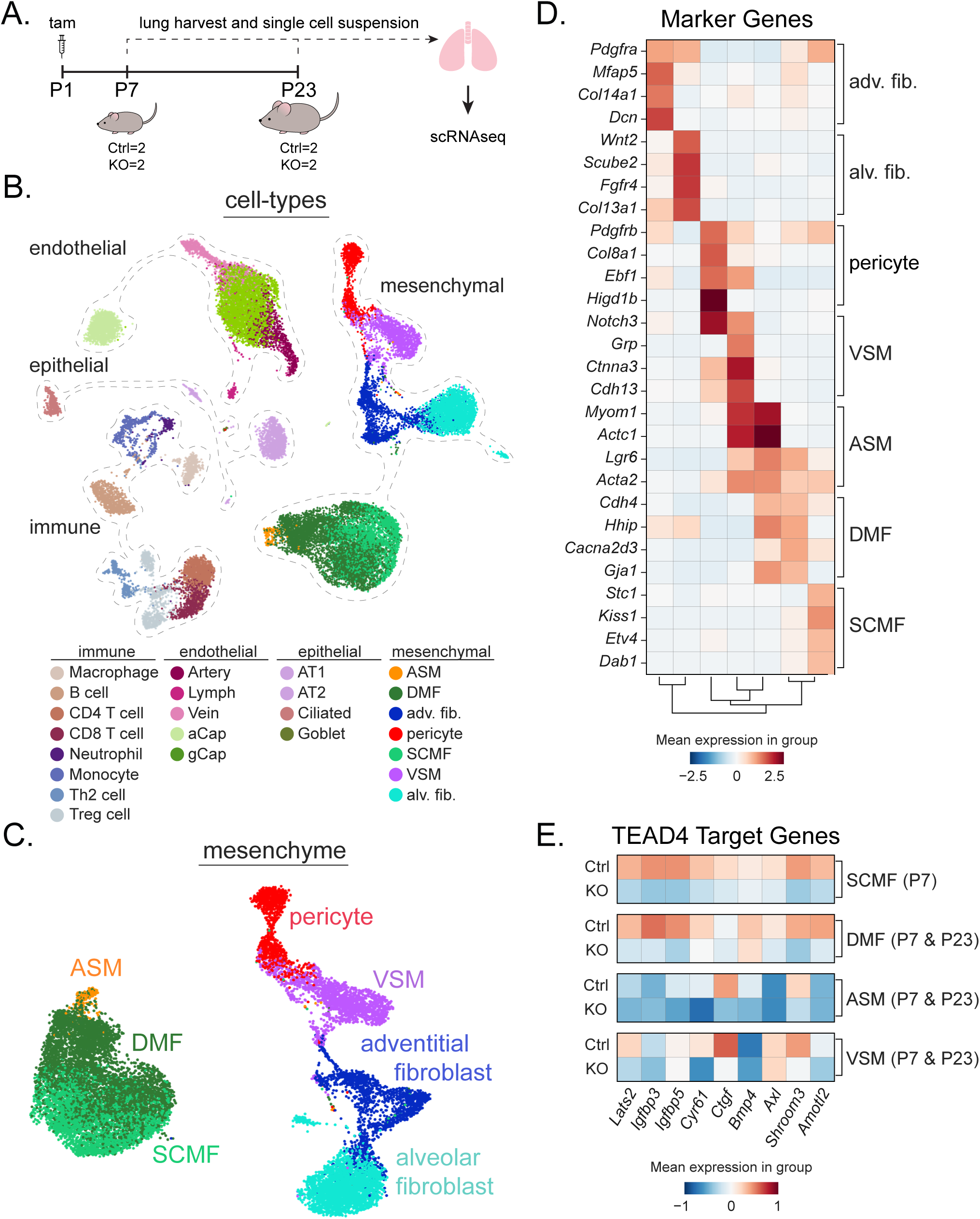
Single-cell RNA sequencing of Ctrl and KO mouse lungs. (A) Experimental schematic of cell isolation from P7 and P23 Ctrl and KO mice and processing for scRNAseq. (B) UMAP of cell types from merged P7 and P23 datasets. The four cell compartments are outlined and each cell type is uniquely colorized and noted in the corresponding legend. (C) UMAP of mesenchymal cell types. (D) Heatmap of marker genes associated with mesenchymal cell types in (C). (E) Heatmap of TEAD4 target gene expression in Acta2-expressing cell types. Expression is consistently reduced in P7 KO SCMFs and DMFs.

We next assessed whether the conditional knockouts had reduced Hippo pathway-regulated genes. Sub-sampling the ASM, DMF, and SCMF from the P7 and P23 timepoints, we filtered our data based on annotated TEAD4 target genes. We found that known Hippo-regulated genes were reduced (**Fig. 4E**). Importantly, the KO SCMFs and DMFs consistently had reduced TEAD4-target gene expression compared to ASM or VSM (**Fig. 4E**). These data show that conditional ablation of Yap/Taz from Acta2-expressing cell types results in impaired expression of known Hippo-pathway target genes.

### ScRNA-seq of Yap/Taz mutant lungs identifies a “ductal myofibroblast-like” population of cells

To determine how the Yap/Taz deletion affected mesenchymal cell populations, we analyzed the subset of *Acta2*-expressing SCMF-related clusters from our scRNA-seq experiments. The UMAP projection of ASM, DMF, and SCMF clusters revealed a unique population of “DMF-like” cells that persisted only in the KO P23 dataset when SCMFs were no longer present (**Fig. 5A**).

**Figure 5.**
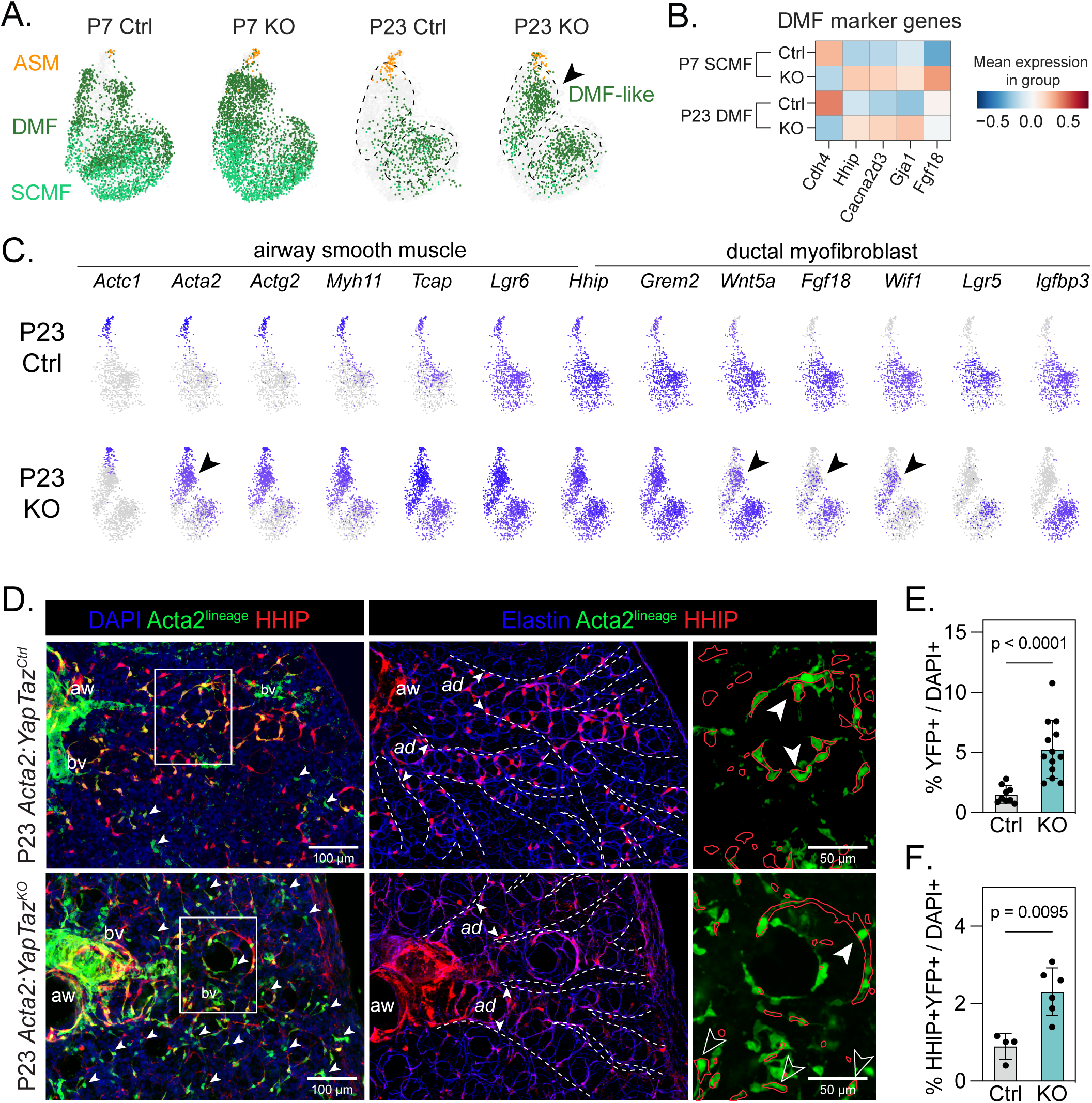
Aberrant expression of HHIP+ DMFs in alveolar ducts of KO mice. (A) UMAP of the SCMF-related ASM/DMF/SCMF cluster, split by age and genotype. Outlined regions highlight a DMF-like population in the P23 KO dataset. (B) Heatmap shows a marked increase in expression of DMF marker genes in both P7 SCMFs and P23 DMFs. (C) Select feature plots show expression of ASM and DMF marker genes in P23 SCMF-related cluster. Arrows emphasize enrichment of smooth muscle markers (*Acta2*) and DMF markers (*Wnt5a*, *Fgf18*, *Wif1*) in the DMF-like KO population. (D) Distal regions of P23 Ctrl and KO PCLS were imaged to capture alveolar ducts. HHIP+ cells mark ductal myofibroblasts and Acta2-lineage traced cells are YFP+. Clusters of YFP+ lineage-traced cells are marked by arrows. Hydrazide was used to mark elastin, which aids in morphologically distinguishing alveolar ducts (a.d., dotted outline). Zoomed in regions highlight aberrant HHIP+ YFP+ myofibroblasts (solid arrow) found in the alveolar ducts of KO mice. Scale bars, 100 µm, 50 µm. (E) Significant increase in the percentage of YFP+ lineage traced cells in P23 KO mice. Ctrl n=8, KO n=11. (F) Increased percentage of HHIP+YFP+ DMFs in P23 KO mice. Ctrl n=4, KO n=6. Data presented as mean ± SD.

A recent study identified the alveolar DMF in scRNA-seq data from a lung developmental time course^21^. The DMF expresses *Hhip* and *Cdh4* marker genes and is spatially distinct from ASM and SCMF^21^. Consistent with the DMF-like population in our scRNA-seq, we observed higher expression of these marker genes in the KO DMF and SCMF cells (**Fig. 5B**). GO term analysis of SCMF-related clusters revealed enriched expression of genes regulating cell population proliferation and differentiation in KO SCMFs (**Fig. S3A**). KO DMFs were enriched in genes associated with negative regulation of cell differentiation (**Fig. S3B**). The GO term analysis indicated that the Yap/Taz deletion may affect changes in cell differentiation and likely the contractile function of *Acta2*-expressing cells, including SCMF.

Further analysis of our SCMF-related clusters at P23 showed a marked increase in *Acta2* expression in our DMF-like KO population. Except for *Actc1*, a committed airway smooth muscle marker, the DMF-like population had higher smooth muscle-associated marker gene expression, largely absent in the Ctrl DMF population. The DMF-like population expressed DMF markers *Wnt5a*, *Fgf18*, and *Wif1*. Expression of Yap target *Igfbp3* marked both Ctrl and KO DMFs. Still, it was notably absent in the *Acta2*-high DMF-like population (**Fig. 5C**). These data indicate that Yap and Taz ablation impairs the differentiation of *Acta2*-expressing DMFs.

### Acta2-lineage-derived cells are increased in the Yap/Taz mutant lungs

While the function of DMF is unknown, it was proposed that DMFs are derived along the same lineage as SCMF but do not undergo apoptosis^21^. To test this in our model, we used the R26R-lox-stop-lox-eYFP (*R26R^eYFP^*) reporter to track Acta2-lineage traced cells in Ctrl and KO conditions. The percent of lineage-traced SCMFs was similar between Ctrl and KO at P7 (∼13% and ∼14%, respectively, data not shown). At P23, the Acta2-lineage+ cells were restricted to the smooth muscle (airway and vascular) and HHIP+ DMFs in Ctrl lungs (**Fig. 5D**). However, in the KO lungs, we found significantly more Acta2-lineage+ cells that persisted at P23 (**Fig. 5D, E**). Furthermore, there were significantly more HHIP+YFP+ cells in the KO lungs (**Fig. 5F**). Imaging revealed irregular HHIP+ DMFs in KO mice. These cells appeared as flattened and elongated and were morphologically distinct from Ctrl DMFs (**Fig. 5D**). These data demonstrate that Yap and Taz in *Acta2*+ cells are required for myofibroblast lineage allocation during alveologenesis.

### Acta2-derived cells persist in the alveolar ducts after alveologenesis

Since we observed a distinct population of cells in the Yap/Taz mutant lungs resembling DMFs, we wanted to confirm that Acta2-lineage tracing during alveologenesis specifically contributes to the DMF cell pool in adult lungs. We induced *Acta2^CreERT2^:R26R^tdTomato^*mice at P7 (neonatal-lin) and harvested tdTomato+ cells well into adulthood (5 months old). We also lineage-traced adult *Acta2*+ cells by inducing adult animals with tamoxifen and tracing for 7 days (adult-lin). We simultaneously harvested lungs from both cohorts and performed scRNA-seq (**Fig. 6A**). The scRNA-seq data from the tdTomato+ cells were integrated with bulk mesenchyme from P7 and Adult scRNA^13^. The tdTomato+ cells from both conditions largely consisted of VSM and a small amount of ASM, which were captured with similar frequency between the two cohorts (**Fig. 6B, C**). Consistent with the hypothesis that the DMF-pool is mostly derived from Acta2+ neonatal cells during alveologenesis, we saw a greater enrichment of the DMF from the neonatal-lin (84%) vs the adult-lin (16%) (**Fig. 6B, C**). Integrating the tdTomato+ neonatal- and adult-lin with published whole P7 and adult mesenchyme showed that SCMFs were completely absent in the adult scRNA-seq (**Fig. 6B**) ^13^. Consistent with the scRNA-seq, imaging revealed an abundance of neonatal-derived tdTomato+ cells expressing HHIP, that resided in alveolar ducts (ad) (**Fig. 6D**). These experiments demonstrate that *Acta2*-expressing neonatal cells persist in the adult lung as DMFs.

**Figure 6.**
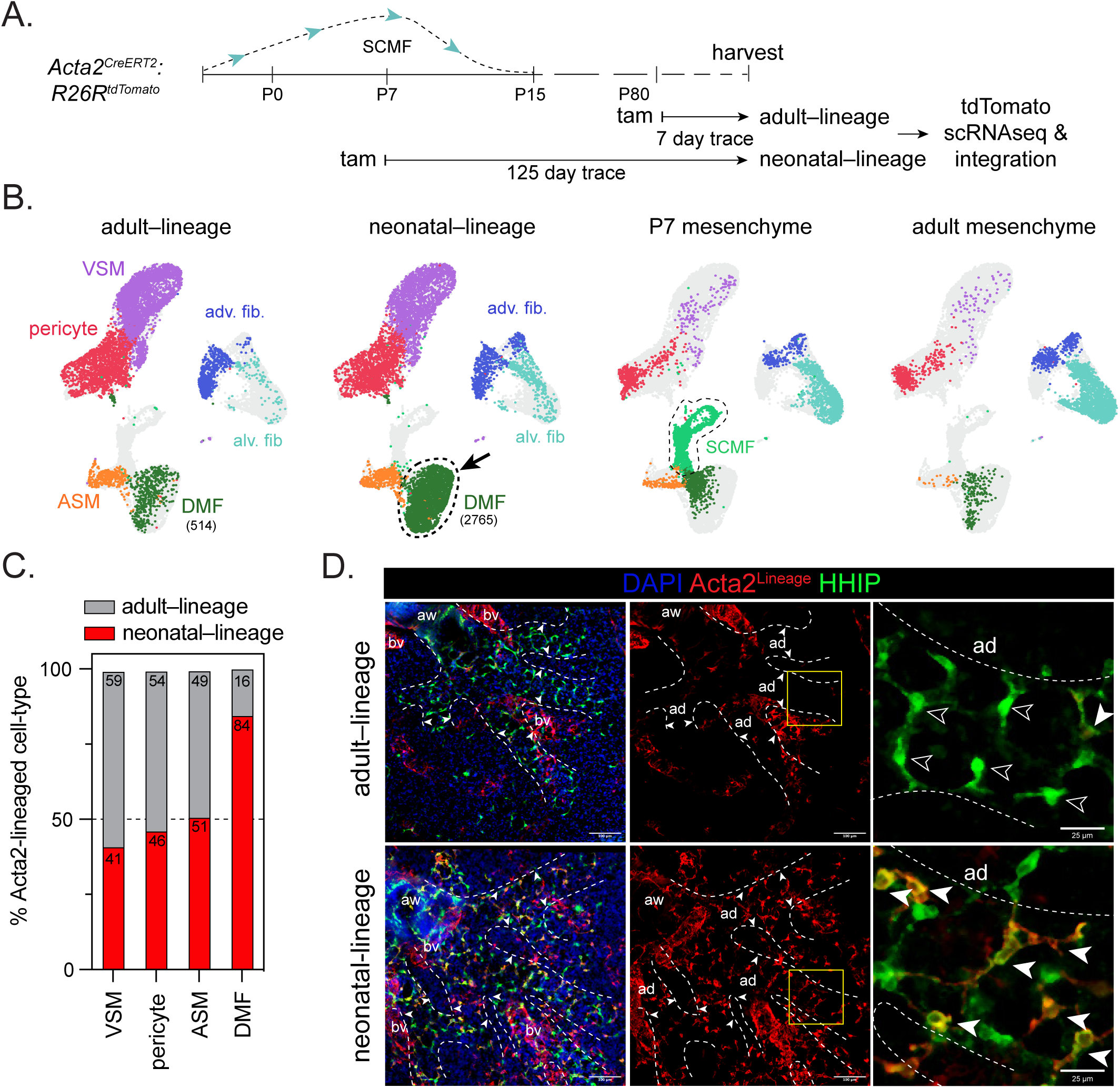
Lineage tracing reveals the SCMF-derived DMF population. (A) Experimental Design: Recombination was induced in *Acta2^CreERT2^:R26R^tdTomato^*mice at P7 and harvested after a 125-day trace (neonatal-lin) or a 7-day trace was done in adult mice (adult-lin). TdTomato+ cells were purified using FACS and scRNA-seq libraries, and sequencing was performed. Datasets were integrated with previously generated P7 and adult mesenchymal datasets. (B) UMAPS of mesenchyme from the indicated experiments, colored by cell type. Notably, there is an increase in the DMF population in the neonatal-lin traced dataset transcriptionally related to the SCMFs at postnatal day 7 (outlined). In paratheses are the “cell” numbers captured in the cell type. (C) Cell-type percentages of Acta2-high cell populations confirm increased contribution to the ductal myofibroblast populations from the neonatal-lin experiment. (D) Imaging of whole-mount tissue with a focus around the distal regions to capture alveolar ducts (a.d, dotted line), shows an increase in lineage traced tdTomato+ cells with HHIP+ DMFs residing in the alveolar ducts in the neonatal-lin lung. Arrows point to HHIP+tdTomato+ cells in the ductal region (outlined). Scale bar, 100 µm, 50 µm.

### Nuclear-Yap induces expression of contractile smooth muscle markers

To assess the role of Yap and Taz in the SCMF lineage, we induced nuclear localization of Yap in cultured neonatal mouse lung fibroblasts. Fibroblasts were seeded in collagen gels and treated with TRULI, a chemical inhibitor of kinases Lats1/2, thereby inhibiting the phosphorylation of Yap (**Fig. 7A**) ^22^. Fibroblasts treated with TRULI expressed higher levels of YAP-TEAD target genes confirming increased Yap nuclear activity (**Fig. 7B**). TRULI treated fibroblasts expressed significantly higher levels of marker genes associated with ASM and DMF. TRULI-treatment significantly reduced the expression of the SCMF marker gene, *Stc1*. However, *Pdgfra* expression was unaffected, suggesting cell-specific Hippo-pathway regulation of target genes **(Fig. 7C)**. Consistent with this, no significant changes were observed in genes encoding markers of other fibroblast populations (**Fig. S1A, B**). Further analysis revealed that neonatal fibroblasts treated with TRULI expressed reduced levels of ECM constituents and markers of apoptosis (**Fig. S1C, D**). These results show that increased YAP-nuclear translocation in neonatal fibroblasts promotes the myofibroblast gene signature and suppresses the SCMF and ECM-related gene program.

**Figure 7.**
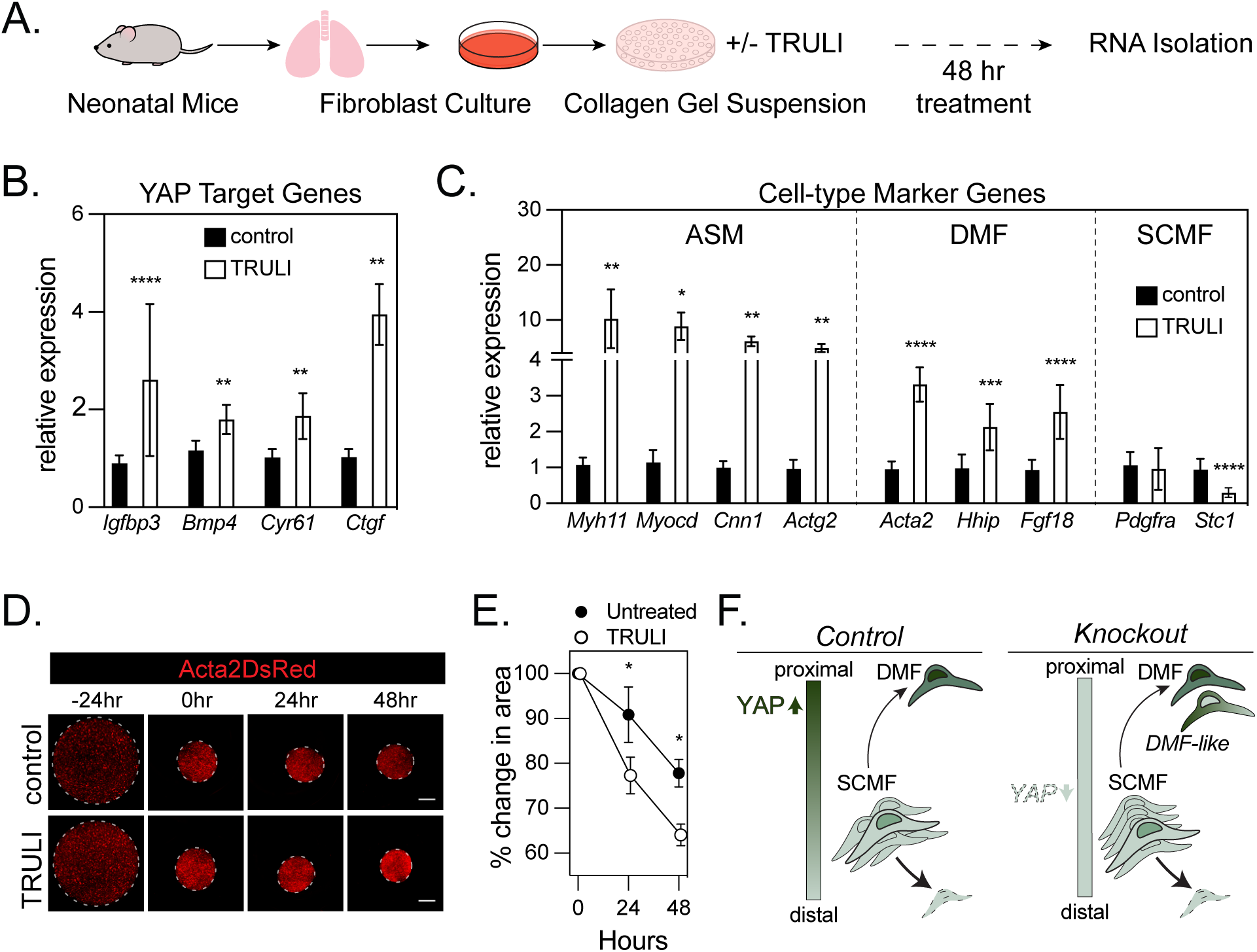
Nuclear-YAP accumulation leads to increased myofibroblast differentiation and function in neonatal fibroblasts. (A) Experimental schematic of lung fibroblast culture from P7 mice; cultured fibroblasts were seeded in collagen gels and treated with TRULI to induce nuclear-YAP, followed by RNA isolation. (B) Increased expression of YAP-TEAD target genes in treated cells confirms TRULI-induced increase in nuclear-YAP. (C) Increased nuclear-YAP leads to an increase in smooth muscle and DMF markers, however, decreases expression of SCMF marker, *Stc1*. Data presented as mean ± SD. (D) Images of collagen gels using primary fibroblasts from postnatal day 7 *Acta2^DsRed^* mice. Cells in collagen gel suspension were treated 24 hours post seeding and the gel area was measured every 24 hours. TRULI treated fibroblasts exhibit increased contraction. (E) Measurements of percent change in collagen gel area support increased contractile activity of TRULI treated fibroblasts. (F) Summary: Increased nuclear-Yap activity promotes differentiation of SCMFs to DMFs and an increased myogenic gene signature. Conditional knockout of Yap/Taz in Acta2-expressing cells resulted in aberrant differentiation of persisting Acta2-lineage traced cells to a contractile DMF-like cell population. Data presented as mean ± SD.

Next, we confirmed that the smooth muscle-associated gene signature had a functional phenotype. We treated fibroblasts from neonatal *Acta2^DsRed^*mice seeded in collagen gels and imaged throughout the experiment (72 hours) to assess their ability to contract the gels (**Fig. 7D**). Consistent with increased expression of contractile markers, cells treated with TRULI had greater DsRed intensity and significantly smaller collagen gels than controls (**Fig. 7E**). These data show that by simulating the Hippo “off” state with TRULI *in vitro*, neonatal fibroblasts assume a contractile smooth muscle over fibroblast phenotype.

Hippo activity increased in transient SCMFs as they underwent developmental apoptosis, evidenced by our bulk sequencing data and a measured decrease in nuclear-YAP expression. scRNAseq and imaging analysis revealed that Acta2-expressing neonatal cells that do not undergo apoptosis, persist as DMFs. Under Yap/Taz knockout conditions, the proximal-distal axis is disrupted, leading to aberrant differentiation. The persisting Acta2-lineage traced cells displayed perturbed differentiation, illustrated by a morphologically and transcriptionally distinct DMF-like cell population (**Fig. 7F**).

## Discussion

Our work provides new insight into the molecular pathways required for the differentiation and function of the neonatal SCMF. We propose that Yap/Taz regulate two crucial aspects of SCMF biology: SCMF maturation/differentiation and function (contractility). Our observations and other recent studies highlight how biomechanical pathways like Hippo can affect cellular development^14, 23–25^. During alveologenesis, we found that dynamic changes in Hippo-pathway occurred in the SCMF. Conditionally knocking out Yap/Taz from Acta2-expressing SCMFs resulted in alveolar simplification. Notably, transcriptional differences between Ctrl and KO conditions were related to cellular differentiation and apoptosis. In line with this, we observed persisting Acta2-lineage traced cells in the KO lungs that expressed similar genes to DMF and ASM. These observations indicate that the Hippo-pathway can alter the balance of mesenchymal differentiation, leading to impaired alveolar maturation. Understanding the origins of mesenchymal cell lineages and underlying pathways that regulate their differentiation could have implications for treating neonatal and adult lung diseases with stromal involvement.

The downstream effectors of the Hippo-pathway, Yap/Taz, relay critical context-dependent signaling that controls cell growth^25^. As alveologenesis proceeds, our data suggests that Yap/Taz regulates the function and differentiation of SCMFs. First, our *in vitro* data indicate that nuclear accumulation of Yap/Taz promotes a myogenic gene program. Using the *Acta2^CreERT2^* line, we consistently observed an alveolar simplification defect in the mutant conditions. A limitation of the current study is that *Acta2* is expressed ubiquitously in the mouse, and we did encounter a colonic obstruction, as previously reported while aging our mice^26, 27^. Further, scRNA-seq identified disrupted gene signatures related to cell differentiation, extracellular matrix production, and proliferation. TEAD4 target genes, including *Igfbp5* and *Shroom3*, were significantly reduced in KO SCMFs. Studies have shown that Igfbp5 modifies ECM and enhances fibrosis, and Shroom3 regulates cell shape through direct interaction with myosin^28–31^. Recently, Mlck and Yap/Taz were ablated from all Pdgfra-expressing fibroblasts, resulting in simplified alveoli, which was only partly due to reduced contractility as demonstrated in phenotypic rescue experiments^12^. Taken together, these data suggest that Yap/Taz regulates a genetic program in addition to the expression of genes required for myogenesis.

In our study, we found that Yap/Taz-mutant lungs had a persisting Acta2-derived cell lineage with transcriptional similarities to the DMF and ASM. The DMFs were recently identified from scRNA-seq and localized in the alveolar ducts^21^. Although the function of these cells is still unknown, it was speculated that DMFs arise from the same progenitors that generate SCMF. Because we observed a persisting cell lineage, we tested this hypothesis and found that, indeed, most of the Acta2-lineaged cells from the neonate that remained in the adult lungs were enriched with DMFs. We localized the Acta2-lineaged cells and found that in KO lungs, the ducts were dilated, and the HHIP+ cells were elongated and more abundant than controls. Although this led us to test whether YAP promoted the differentiation of DMFs, the abundance of these cells in the KO may have non-cell autonomous effects on alveolar maturation. These observations warrant further studies to target these cells using genetic tools and test their function.

We observed that the deletion of Yap/Taz affects the differentiation of SCMFs. We noted a decrease in YAP nuclear localization in SCMFs by P7 and P10, coinciding with the initial appearance and expansion of SCMFs. Although SCMFs are normally programmed to undergo apoptosis, conditionally removing YAP resulted in aberrant differentiation. More detailed temporal experiments are needed to determine whether intrinsic or extrinsic apoptotic pathways are activated in SCMF. In Drosophila models of Hippo pathway signaling, Yap/Taz is generally considered pro-growth/proliferation^32, 33^. However, in mammalian systems, the crosstalk with additional developmental pathways and cell-type specificity adds complexity that may have arisen through evolution. Studies have shown that in the presence or absence of additional pathways, Hippo-YAP signaling is pro-apoptotic. For example, *in vitro*, YAP can interact with p73 and promote intrinsic apoptosis due to DNA damage^34^. Further work *in vivo* suggests that in adult hepatocytes, high levels of Yap during proliferation are balanced by increased susceptibility to apoptosis^35, 36^. In this setting, Yap-mediated cell growth requires a second extrinsic signal to overcome apoptosis in favor of proliferative expansion^36^. In the lung, signaling pathways, including Pdgf, Shh, Tgfb, and Wnt, are required for alveologenesis and affect different aspects of mesenchymal differentiation^6, 11, 13, 37–43^. In all, these ligands may provide the additional signals that inform the balance of proliferation or apoptosis, while the Hippo-pathway relays biomechanical cues along the proximal-distal axis.

The mechanisms of Yap/Taz regulating SCMF differentiation and function may have implications for understanding human lung diseases. Dynamic YAP nuclear localization, normally required for SCMF maturation, proliferation, and apoptosis, could be disrupted in preterm infants with disrupted alveologenesis. Consistent with this, one might expect that the DMF, which resides in the alveolar duct, would also be affected in neonatal lung diseases. The alveolar ducts in mice are structurally similar to the small airways or respiratory bronchioles in humans, which may have implications for small airway disease and adult chronic obstructive pulmonary disease (COPD) ^44–46^. Our work suggests that there exists a proximal to distal gradient of Yap nuclear localization in ASM, DMF, and SCMF, respectively. These observations require further experiments to characterize YAP localization along this axis. Nevertheless, the SCMF is a distinctive mesenchymal cell in the lung. Identifying the molecular regulators like Yap/Taz that underlie SCMF expansion, differentiation, and apoptosis after completing their critical role in alveolar development will shed light on strategies for cell-targeted therapies.

## Acknowledgments

We thank Florin Tulic and the CHOP Flow Cytometry Core; Teodora Orendovici and the CHOP High-Throughput Sequencing Core; and the Center for Applied Genomics at CHOP. We are grateful for the generous support provided by the Ayla Gunner Prushansky Research Fund. This work was supported by the National Institutes of Health (R00HL141684, R35GM119461 to J.A.Z.; R01HL119503, K24HL143281 to L.R.Y.) and the Rare Lung Diseases Frontier Program, Children’s Hospital of Philadelphia to L.R.Y, S.M.M., J.A.Z.

## Methods

### Mice

All animal procedures and experiments complied with guidelines and protocols reviewed and approved by The Children’s Hospital of Philadelphia Institutional Animal Care and Use Committee. The *Acta2^DsRed^* reporter mouse was generously provided by Dr. David Brenner^47^. The *Acta2^CreERT2^* mouse strain was generously provided by Dr. Pierre Chambon and Dr. Daniel Metzger^48^. The following strains were obtained from Jackson Laboratories: *PdgfraH2B^:EGFP^*(Stock no. 007669), *Rosa26R^eYFP^* (Stock no. 006148), *Taz^Flox^/Yap^Flo^*^x^ (Stock No. 030532), *Rosa26R^tdTomato^* (Stock no. 007914). All mice were maintained on an outbred CD-1 background (Charles River Laboratories). *Acta2^CreERT2^:Rosa26R^eYFP^*mice were crossed with *Taz^Flox^/Yap^Flo^*^x^ mice. All experiments used littermates, and male and female mice were analyzed. Experimental mice that are referred to as *Acta2^CreERT2^:Yap/Taz^Ctrl^*(also referred to as “Ctrl”) are *Acta2^CreERT2^:Yap^F/+^:Taz^F/+^,* and *Acta2^CreERT2^:Yap/Taz^KO^* (referred to as “KO”) are *Acta2^CreERT2^:Yap^F/F^:Taz^F/F^*.

### Tamoxifen induction of cell-lineage tracing

Tamoxifen (cat. # T006000, Toronto Research Chemicals) was dissolved in corn oil (cat. # C8267, Sigma-Aldrich) at a concentration of 20 mg/ml. For early postnatal induction at P1, or neonatal induction at P7, tamoxifen (200 mg/kg) was administered via intraperitoneal injection using a 30G needle, after which the injection site was pinched for ten seconds to prevent tamoxifen leak from the peritoneal cavity. Adult animals were given tamoxifen (200 mg/kg) via oral gavage. P1 induced Ctrl and KO experimental mice were harvested at P7 and P23. *Acta2^CreERT2^:R26R^tdTomato^*neonatal mice induced at P7 were harvested after a 125-day trace and are referred to as “neonatal-lin”. *Acta2^CreERT2^:R26R^tdTomato^* adult mice were induced at 12 weeks of age and harvested after a 7-day trace and are referred to as “adult-lin”.

### Histology and Immunofluorescence

Mice were euthanized by CO2 inhalation. After exposing the chest cavity, the lung was cleared of blood by perfusing into the right ventricle with cold PBS. The lungs were inflated with 2% or 4% paraformaldehyde (PFA) under a constant pressure of 15 to 30 cm water and allowed to fix overnight at 4°C.

For staining and immunohistochemistry (IHC), the tissue was gradually dehydrated to 100% ethanol and paraffin-embedded for sectioning. Sections were stained with H&E and Masson’s trichome. For IHC, heat-induced antigen retrieval was performed for 30 seconds at 120°C, followed by 10 seconds at 90°C in a Decloaking Chamber (BioCare Medical) in Reveal Decloaker (cat. # MSPP-RV1000M, Biocare Medical), followed by a brief incubation in 3% H_2_O_2_. Samples were blocked using 5% donkey serum and incubated with primary antibodies in 0.1% PBS-T overnight at 4°C. Primary antibody amplification was performed using ImmPRESS HRP Polymer Detection Kit (cat. # MP-7402-50, Vector Laboratories) and TSA TMR Reagent (cat. # SAT702001, Akoya Biosciences). The following antibodies were used: rabbit anti-proSP-C (1:100, cat. # AB3786, Chemicon), mouse anti-HOP (1:100 with amplification, cat. # sc-398703, Santa Cruz), rabbit anti-YAP (1:100 with amplification, cat. # 4912, Cell Signaling), guinea pig anti-RFP (1:100, cat. # 390 004, Synaptic Systems).

Sectioning and staining for precision-cut lung slices (PCLS) were carried out as follows. After overnight fixation in 2 or 4% PFA, the lung tissue was washed three times with cold PBS. Lungs were cut into smaller transverse sections before embedding in 4% low melting agarose (cat. # IB70050, IBI Scientific). The embedded tissue was sectioned at 150-200 µm thickness using a Leica Vibratome (VT1000S). Sections were permeabilized using perm-buffer, PBS + 1% Triton-X100 (cat. # T8787, Sigma-Aldrich), for one hour at room temperature. Tissue sections were stained with primary antibodies in blocking buffer (PBS + 0.3% Triton-X100 + 1% BSA (cat. # B2518, Sigma-Aldrich)) for 48 hours on a rocker at 4°C. The tissue was washed with perm-buffer three times at room temperature for 30-minute incubations. The tissue and secondary antibodies were incubated for 24 hours at 4°C on a rocker in blocking buffer. The sections were further cleared with Scale A2 and Scale B4 for one week each before imaging by confocal microscopy as described previously^13^. The following antibodies were used: rabbit anti-TAGLN (1:1000, cat. # AB14106, Abcam), chicken anti-GFP (1:500, cat. # GFP-1020, Aves Labs), goat anti-HHIP (1:500, cat. # AF1568, R&D Systems), rabbit anti-ERG (1:500, cat. # ab92513, Abcam), rabbit anti-RFP (1:500, cat. # 600-401-379, Rockland Immunochemicals). Alexa Fluor 647 Hydrazide (1:500, cat. # A20502, Invitrogen) was used to label elastin fibers^17^.

The following secondary antibodies were used at a dilution of 1:500: donkey anti-rabbit IgG Alexa Fluor 555 (cat. # A32794, Invitrogen), donkey anti-rabbit IgG Alexa Fluor 647 (cat. # A32795, Invitrogen), donkey anti-goat IgG Alexa Fluor 488 (cat. # A32814, Invitrogen), donkey anti-goat IgG Alexa Fluor 555 (cat. # A32816, Invitrogen), donkey anti-guinea pig IgG Alexa Fluor 488 (cat. # 706-545-148, Jackson ImmunoResearch), donkey anti-chicken IgG Alexa Fluor 488 (cat. # 703-545-155, Jackson ImmunoResearch). DAPI (2.5 µg/ml, cat. # D9542, Sigma) was used to stain nuclei.

### Imaging and Quantification

Images were obtained with a Leica SP8 confocal microscope or Leica DMi8 Thunder microscope system. Image processing was performed in ImageJ (FIJI). The PCLS images are presented as z-stack projections of 100-150 µm imaged by optical slices of 1 µm.

Paraffin-embedded sections from *Acta2^DsRed^* mice were stained with YAP antibody. Representative images in Figure 1E have been pseudo-colored. Using the spot detection and colocalization analysis tool in Imaris, total nuclear YAP in alveolar DsRed+ fibroblasts was quantified. Paraffin-embedded sections from *Acta2^Ctrl^* and *Acta2^KO^*mice were stained with HOPX and SPC to mark AT1 and AT2 cells, respectively. The spot detection tool in Imaris was used to quantify total HOPX and SPC cells. PCLS images were used to quantify alveolar diameters which were measured using the line tool in ImageJ (FIJI). Background stains such as hydrazide were used to demarcate alveoli and measurements were collected at 20 µm intervals through a z-stack of at least 125 µm. SM22+ alveolar fibroblasts and Acta2+ lineage traced cells were quantified using the spot detection tool in Imaris. Images of distal and proximal alveolar regions from PCLS tissue with endogenous YFP and HHIP staining were used for quantification. Hydrazide 647 has been pseudo-colored to blue in Figure 5D for enhanced contrast. The Imaris colocalization analysis tool was used to find YFP+ HHIP+ myofibroblasts. The spot detection tool was used to quantify total HHIP+ ductal myofibroblasts, YFP+ lineage traced myofibroblasts, and YFP+ HHIP+ myofibroblasts. When quantifying alveolar fibroblasts, thresholding was adjusted manually, and spot removal was used to ensure airway and vascular smooth muscle cells were not quantified.

### Lung Digestion for single-cell suspension

Lung tissue was minced with a razor blade and resuspended in a digestion buffer containing of 480 U/ml Collagenase Type I (Gibco), 50 U/ml Dispase (Corning), and 0.33 U/ml DNase (Roche). The digestion mix was incubated at 37°C for 20-45 minutes, with frequent agitation. The cell solution was filtered through a 100 µm cell strainer. ACK lysis buffer was used to remove red blood cells, followed by 40 µm filtration. The cell pellet was resuspended in FACS buffer (HBSS + 2% FBS + 25 µM HEPES +2 mM EDTA).

### Cell Culture

Primary mouse lung fibroblasts were isolated from early neonatal mice following the single cell suspension prep above. Cells were cultured for no more than 4 passages prior to use in assay. Cells were cultured in DMEM/F-12 (cat. # 11320033, Gibco), were washed using DPBS (cat. # 21-031-CV, Corning), and 0.05% Trypsin-EDTA (cat. # 25300054, Gibco) was used for cell dissociation.

Primary fibroblasts were resuspended in serum-free DMEM/F-12 at 5x10^5^ cells/ml for the collagen gel contraction assay. Next, fibroblasts were mixed with Cultrex rat tail collagen 1 (cat. # 3447-020-01, R&D Systems, 5mg/ml) for a final 1mg/ml collagen concentration. NaOH (cat. # S2770, Sigma-Aldrich) was added to the cells/gel mix to obtain a pH of ∼ 7.5, and gels were dispensed in a 48-well plate. Collagen gels were allowed to solidify for 1 hour at 37°C and were detached from the walls of the well by running a 10 µl pipette tip around the periphery. Collagen gels were resuspended in low serum media (DMEM/F-12 + 2% FBS) and allowed to contract for 24 hours before treatment. After 24 hours, TRULI (2.5 µM, cat. # E1061, Selleck Chemicals) was added to gels in low serum media. Collagen gels were imaged using Leica DMI8 Thunder microscope at 24- and 48-hours post-treatment. The area of gels was measured using the elliptical selection tool in ImageJ (FIJI). These data are presented as percent change in area 24- and 48-hours post-treatment. The assay was completed a minimum of 3 times and the presented data is one from representative experiment.

### RNA Isolation and qPCR

RNA was isolated from cells collected in Trizol (cat. # 15596018, Invitrogen) according to the manufacturer’s protocol. The cDNA was generated from purified RNA using the SuperScript III First-Strand Synthesis System (cat. # 12574026, Thermo Fisher), and real-time PCR was performed using Power SYBR Green reagents (cat. # 4367659, Thermo Fisher) on a QuantStudio 5 Real-Time PCR System (Applied Biosystems).

The following primer sequences were used: *Gapdh,* for: aggtcggtgtgaacggatttg, rev: ggggtcgttgatggcaaca, *Acta2*, for: cccagacatcagggagtaatgg, rev: tctatcggatacttcagcgtca, *Fgf18,* for: cctgcacttgcctgtgtttac, rev: tgcttccgactcacatcatct, *Hhip,* for: gaagatgctctcgtttaagctgc, rev: ccaccacacaggatctctcc, *Myh11,* for: atgaggtggtcgtggagttg, rev: gcctgagaagtatcgctccc, *Myocd,* for: cagtgaagcagcaaatgactcgg, rev: gtcgttggcgtagtgatcgaagg, *Cnn1*, for: tctgcacattttaaccgaggtc, rev: gccagcttgttctttacttcagc, *Actg2,* for: ccgccctagacatcagggt, rev: tcttctggtgctactcgaagc, *Igfbp3,* for: cacaccgagtgaccgattcc, rev: gtgtctgtgctttgagactcat, *Bmp4,* for: attcctggtaaccgaatgctg, rev: ccggtctcaggtatcaaactagc, *Cyr6,* for: taaggtctgcgctaaacaactc, rev: cagatccctttcagagcggt, *Ctgf*, for: ggcctcttctgcgatttcg, rev: gcagcttgacccttctcgg, *Casp3,* for: tggtgatgaaggggtcatttatg, rev: ttcggctttccagtcagactc, *Cdkn1a,* for: cctggtgatgtccgacctg, rev: ccatgagcgcatcgcaatc, *Bak1*, for: caaccccgagatggacaactt, rev: cgtagcgccggttaatatcat, *Bax*, for: tgaagacaggggcctttttg, rev: aattcgccggagacactcg, *Stc1*, for: acgaggcggaacaaaatgatt, rev: tgcactttaagctctctttgaca, *Pdgfra,* for: tatcctcccaaacgagaatgaga, rev: gtggttgtagcaagtgtacc, *Dcn,* for: tcttgggctggaccatttgaa, rev: catcggtaggggcacataga, *Col14a1*, for: tttggcggctgcttgtttc, rev: cgcttttgttgcagtgttctg, *Mfap5,* for: tcaacgcggagatgatgtgc, rev: tcagccagagctgtatcgtct, *Wnt2a*, for: tctgaaacaagaatgcaagtgtca, rev: gagatagtcgcctgttttcctgaa, *Fgfr4,* for: aggagccaggaaggtggtca, rev: aaagatgctcaacagggccaa, *Scube2*, for: ccatgctgatgcactgtgtca, rev: agtgtgttgtcacattcatccat, *Pdgfrb*, for: aggagtgataccagctttagtcc, rev: ccgagcaggtcagaacaaagg, *Notch3*, for: agtgccgatctggtacaactt, rev: cactacggggttctcacaca, *Eln*, for: ttgctgatcctcttgctcaac, rev: gcccctggataatagactccac.

### Bulk RNA Sequencing

Lungs from *Acta2^DsRed^:Pdgfra^GFP^* mice at the indicated timepoints were processed to generate a single-cell suspension as described above. Cells were negatively sorted for DAPI, and the RFP+ GFP+ double positive population was collected. The SCMFs were recovered in FACS buffer, centrifuged, and the cell pellet was lysed with 1 ml Trizol. The RNA was isolated, and libraries were prepared as described previously with the following modifications: Libraries were prepared using the SMART-seq v4 Plus/ultra-low RNA input stranded total RNA-seq kit (Takara Bio Inc.) and sequenced on the NextSeq 1000/2000 (100 cycles) v3 platform (Illumina) ^8^.

Reads in FASTQ format were aligned to the NCBI RefSeq mouse reference genome (mm10/GRCm38) and quantified at the gene level using the nf-core/rnaseq v3.8.1 Nextflow pipeline^49, 50^. Differential gene expression analysis was performed using DESeq2^51^. Differentially expressed genes (DEGs) were identified between groups using the *ashr* results test, and were called with a false-discovery rate (FDR) cutoff of FDR < 0.05 and a minimum log2-fold change (LFC) of magnitude 0.25^52^. Variance stabilized counts used for plotting and tertiary analysis were generated using the *rlog* function with *blind=FALSE*. Tertiary processing of bulk-seq data was performed using scanpy^53^. Gene set scoring was performed using the JASMINE scoring metric^54^.

### Single Cell RNA Sequencing

For single-cell RNA sequencing, cells were negatively sorted for DAPI and both CD45 (APC, cat. # 17-0451-82, eBioscience) negative and positive populations were collected in FACS buffer without EDTA using the MoFlo Astrios Cell Sorter. Cells were pelleted and counted by trypan blue exclusion and CD45- and CD45+ cells were combined at a 5:1 ratio and resuspended according to the 10X Genomics protocol, aiming for 10,000 targeted cell recovery. The single-cell suspension was loaded onto the 10X Chromium Controller and libraries were prepared according to the Chromium Next GEM Single Cell 3’ v3.1 Dual Index Library protocol.

Reads in FASTQ format were aligned to the NCBI RefSeq GRCm39 genome, and reads with unique molecular identifiers (UMIs) were quantified at the gene level using STAR-solo, with *soloUMIdedup=”1MM_CR”*, *soloUMIfiltering=”MultiGeneUMI_CR”*, and *soloCellFilter=”EmptyDrops_CR”* ^55, 56^. Ambient RNA was removed from the generated counts matrices using the scvi-tools implementation of scAR^57, 58^. Ambient-corrected counts matrices were subsequently processed using scanpy and anndata^53, 59^.

Cells were filtered to include only those cells with more than 1000 unique genes detected/cell, fewer than 6049 unique genes detected/cell (the 95th percentile), and fewer than 15% of UMIs attributed to mitochondrial genes. Genes were filtered from the combined dataset if detected in fewer than 47 cells (0.1% of total number of unfiltered cells). Putative cell doublets were identified and removed from each batch independently using the scvi-tools implementation of the SOLO doublet identification model^60^.

Gene expression was calculated from scAR-corrected UMI-counts by first normalizing to CPM using the *sc.pp.normalize_total* function with *target_sum=1e6* and then calculating ln(CPM + 1) using the *sc.pp.log1p* function. The top 2000 highly variable genes in the dataset were identified using the *sc.pp.highly_variable_genes* function, with *flavor=”seurat_v3”* and *n_top_genes=2000*. An initial principal component analysis was performed using the *sc.tl.pca* function with *svd_solver=”arpack”*. This PCA was subsequently batch-corrected using the pytorch implementation of the harmony batch correction algorithm^61^. A cell-cell neighborhood was calculated using the *sc.pp.neighbors* function, with n_neighbors=*int(0.25 * sqrt(adata.n_obs)),* and this cell-cell neighborhood was used to generate a 2-dimensional UMAP using the *sc.tl.umap* function with *init_pos=”spectral”* ^62^. Finally, cell clustering was performed using the leiden algorithm via the *sc.tl.leiden* function with *resolution=1*^63^.

Putative cell compartment (and subsequently cell type) labels were assigned to cells using the scvi-tools implementation of the SCANVI model^64^. Briefly, a small set of cells were used as ‘seeds’ for each of the expected cell compartment/type categories, identified by their maximal expression of established marker genes for each cell compartment/type. The gene expression profiles of these ‘seed’ cells were then used to build the SCANVI model and predict putative cell compartment/type labels of unlabeled cells in the dataset. After this labeling step, cell compartment/type labels were collapsed on a per-cell-cluster (leiden cluster) basis to refine the final cell compartment/type labels using the additional information provided by cell-cell neighborhood network clustering.

### GO Term Volcano Plots

Over-representation analysis (ORA) was performed using gProfiler, with runtime options *unordered query*, *only annotated genes*, *g:SCS threshold < 0.05*, and *max term size = 2500*. Biological processes were filtered to exclude duplicate terms between control and knockout datasets. Pathways and processes were selected based on relevance to lung development and terms specifying directionality^65^.

### Statistics and Reproducibility

All statistical analyses were performed in GraphPad Prism 9.0 using two-tailed Student’s t-tests, unless otherwise indicated in figure legends. Differences were considered significant if the p-value was < 0.05. All animal studies are from multiple litters and quantification data presented are averaged values from individual mice where indicated. All *in vitro* experiments were performed at least three times with successful replication. Specific n values are listed in the figure legends.

