## Supplementary figures and images for "Dynamic Hippo pathway activity underlies mesenchymal differentiation during lung alveolar morphogenesis"

### Suppl. Figure 1

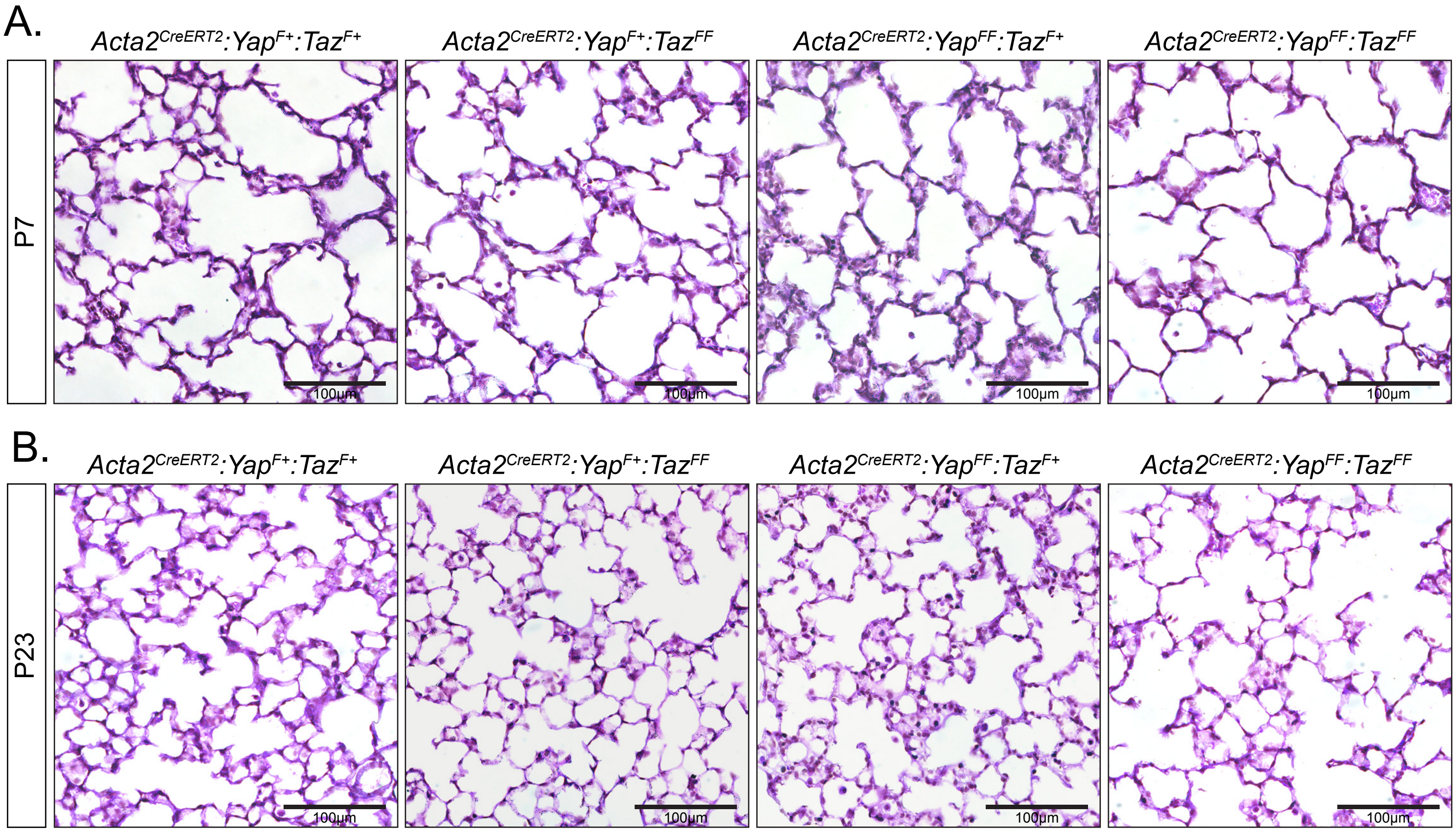

### Suppl. Figure 2

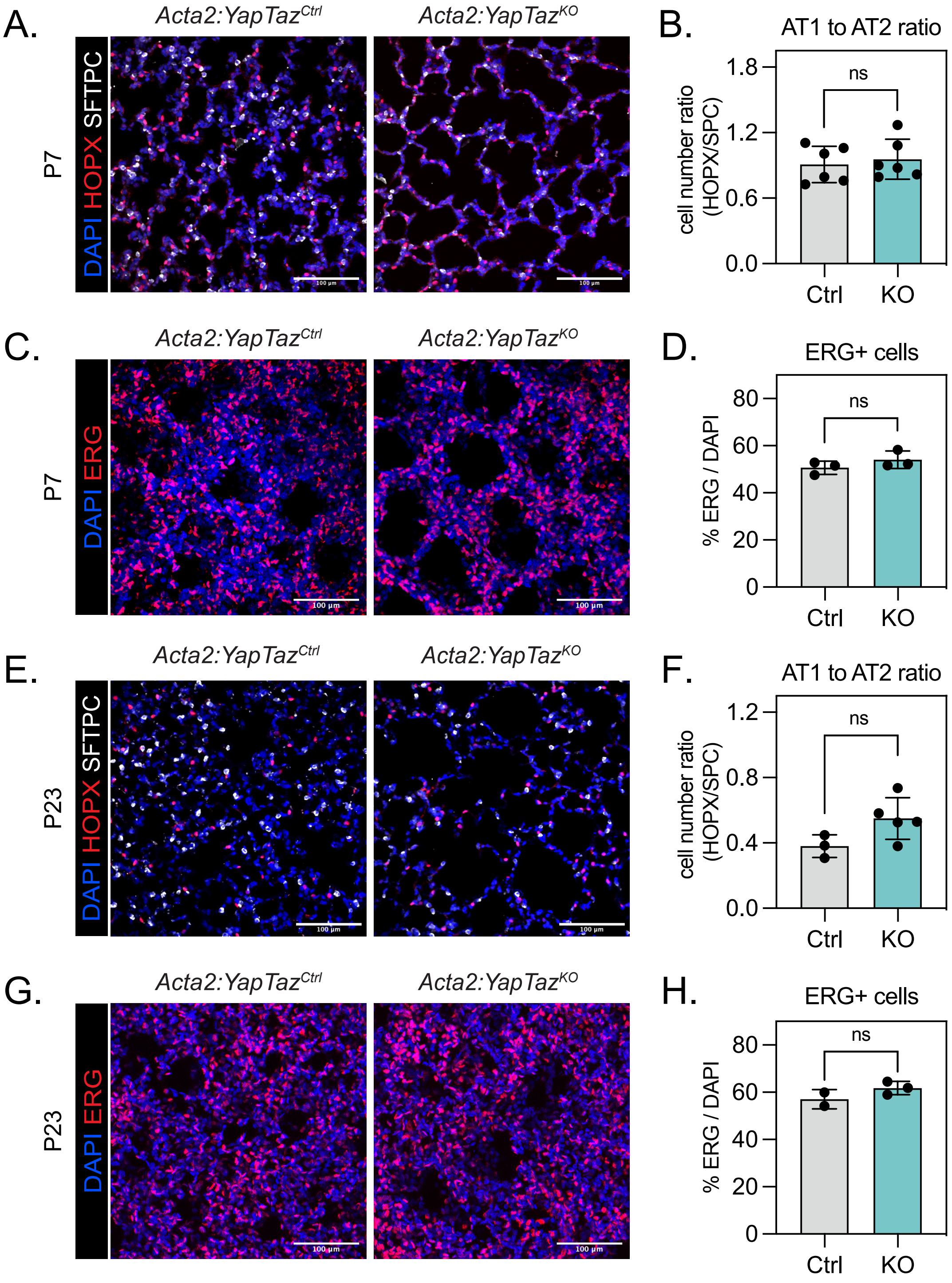

### Suppl. Figure 4

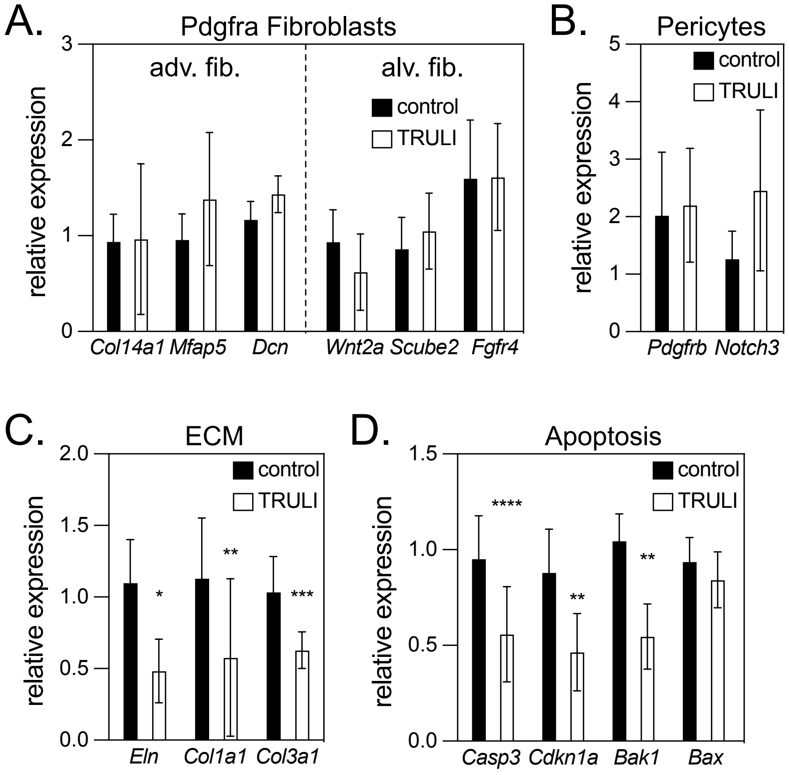
