## Supplementary material for "Dynamic Hippo pathway activity underlies mesenchymal differentiation during lung alveolar morphogenesis": Suppl. Figure 3

**Figure S3.**

A.

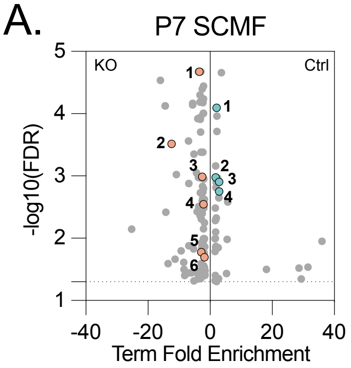


P7 Ctrl vs. KO SCMF : GO:BP Term

reduced in KO
(1) tube development
(2) cell migration
(3) response to decreased oxygen levels
(4) response to hypoxia

higher in KO
(1) cell morphogenesis involved in differentiation
(2) semaphorin-plexin signaling pathway
**(3) positive regulation of cell differentiation**
**(4) positive regulation of cell population proliferation**
(5) regulation of apoptotic process
(6) positive regulation of cell migration

B.

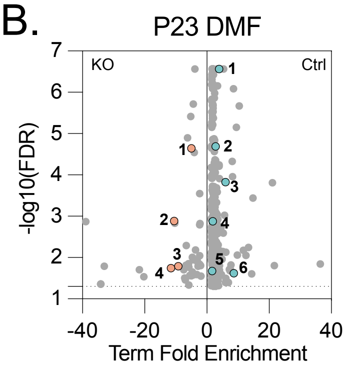


P23 Ctrl vs KO DMF : GO:BP Term

reduced in KO
(1) extracellular matrix organization
**(2) positive regulation of cell differentiation**
(3) positive regulation of smooth muscle cell migration
(4) Wnt signaling pathway
**(5) negative regulation of cell population proliferation**
(6) positive regulation of Notch signaling pathway

higher in KO
**(1) muscle contraction**
(2) myofibril assembly
**(3) negative regulation of muscle cell differentiation**
(4) mesenchyme morphogenesis
