## Supplementary material for "Dynamic Hippo pathway activity underlies mesenchymal differentiation during lung alveolar morphogenesis": Suppl. Figure Legends

**Supplemental Figure Legends**

**Figure S1. Conditional knockout of either Yap or Taz does not contribute to alveolar simplification.** (A) Recombination was induced at P1 and lungs were harvested at P7. H&E staining of mouse lungs at P7 displaying differences in alveolar architecture between *Acta2^YapF+/TazF+^* (Ctrl), *Acta2^YapF+/TazFF^* (TazKO), *Acta2^YapFF/TazF+^* (YapKO), and *Acta2^YapFF/TazFF^* (KO). Ctrl n=9, TazKO n=4, YapKO n=10, KO n=8. (B) Lungs from mice induced at P1 were analyzed at P23. H&E staining of P23 mouse lungs shows striking alveolar simplification of KO lung compared to Ctrl and single allele knockouts. Ctrl n=8, TazKO n=8, YapKO n=6, KO n=10.

**Figure S2. Epithelial and endothelial numbers are unaffected in KO conditions.** Tamoxifen was administered to neonatal pups at P1 and analyzed at the indicated timepoints. (A) Immunostaining of AT1 (HOPX) and AT2 (SFTPC) cells in the alveolar space of Ctrl and KO mice at P7. Scale bar, 100 µm. (B) Quantification of AT1 (HOPX+) to AT2 (SFTPC+) cell number ratio between P7 Ctrl and KO mice. Ctrl n=6, KO n=6. (C) Whole mount imaging of ERG staining on Ctrl and KO P7 tissue. Scale bar, 100 µm. (D) Quantification of ERG+ cells in alveolar regions. Ctrl n=3, KO n=3. (E) Immunostaining of AT1 (HOPX) and AT2 (SFTPC) cells in the alveolar space of Ctrl and KO mice at P23. Scale bar, 100 µm. (F) Quantification of AT1 (HOPX+) to AT2 (SFTPC+) cell number ratio between P23 Ctrl and KO mice. Ctrl n=3, KO n=5. (G) Whole mount imaging of ERG staining on Ctrl and KO P23 tissue. Scale bar, 100 µm. (H) Quantification of ERG+ cells in alveolar regions. Ctrl n=2, KO n=3. Data presented as mean ± SD.

**Figure S3. Biological processes associated with proliferation and differentiation impacted by *Yap/Taz* deletion in *Acta2*+ cells.** The indicated cell-types were sub-selected from scRNA-seq, differential gene expression was calculated, and GO:BP-term analyses were performed. Numbered colorized points on volcano plot correspond to the accompanying list of select GO Terms with a focus on pathways or processes such as proliferation and differentiation (bold) (A) Volcano plot of GO:BP term analysis for P7 SCMFs. Upregulated genes in KO SCMFs are enriched in terms associated with positive regulation of differentiation and proliferation.(b) Volcano plot of GO:BP term analysis for P23 DMFs. Upregulated genes in KO DMFs are enriched in terms associated with muscle contraction and negative regulation of cell differentiation.

**Figure S4. Increased nuclear-Yap attenuates expression of markers for apoptosis and ECM components.** (A, B) Expression of *Pdgfra+* adventitial and alveolar fibroblast markers and pericyte markers are unaffected by TRULI treatment. (C) TRULI treatment in neonatal fibroblasts reduced expression of ECM components. (D) Markers of apoptosis are reduced in TRULI treated conditions. Data presented as mean ± SD.
